# An evolutionarily conserved metallophosphodiesterase is a determinant of lifespan in *Drosophila*

**DOI:** 10.1101/2020.05.08.084137

**Authors:** Kriti Gupta, Vishnu Janardan, Sanghita Banerjee, Sveta Chakrabarti, Swarna Srinivas, Deepthi Mahishi, Padinjat Raghu, Sandhya S. Visweswariah

## Abstract

Evolutionarily conserved genes usually have a critical role to play during organismal aging and longevity. Here, we show that a previously uncharacterized Class III metallophosphoesterase in *Drosophila*, an ortholog of the MPPED1 and MPPED2 proteins in mammals, is necessary for optimal lifespan. dMPPED is the product of the gene *CG16717* and hydrolyzed a variety of phosphodiester substrates in a metal-dependent manner. *dMPPED* was expressed widely during development and in the adult fly. Deletion of the gene in flies dramatically reduced lifespan, without affecting development or fecundity. Longevity was restored on ubiquitous expression of the protein, and neuronal expression of both wild type and the catalytically inactive form of dMPPED was also able to restore normal lifespan. Overexpression of the protein, both ubiquitously and neuronally in wild type flies extended lifespan by ~ 20%. RNA-seq analysis of *dMPPED*^*KO*^ flies revealed mis-regulation of innate immune pathways, a number of transcription factors and genes earlier reported to affect aging and lifespan. Importantly, neuronal expression of mammalian MPPED2 was able to rescue lifespan in *dMPPED*^*KO*^ flies, but not extend lifespan in wild type flies. This reports the first description of the biological role of an evolutionarily conserved metallophosphoesterase that may serve as a scaffolding protein in diverse signaling pathways to modulate longevity in the fly.

## Introduction

Proteins that are evolutionarily conserved usually play a role in fundamental biological processes in an organism [1, 2]. Sequence conservation at the level of amino acids, especially at the catalytic site of an enzyme, implies strong structural conservation and perhaps common activities, in organisms separated by large tracts of time. If a gene linked to a genetic disease in humans has a counterpart in lower organisms more amenable to genetic manipulation, important insights into the gene’s function can be gained by either deletion or overexpression of the ortholog in a simpler organism. Such studies pave the way towards defining approaches that can be utilized in mammalian model systems at a later date.

WAGR syndrome (Wilms’ tumor, aniridia, genitourinary anomalies, mental retardation) [3] is associated with interstitial deletions in a region in chromosome 11p13 [4, 5]. This locus contains a number of genes such as *WT1, BDNF* [6], *PAX6* [7] (all important during development), *FSHB* [8], *RCN1*, and *MPPED2* (*239FB*) [9, 10]. *MPPED2* mRNA is predominantly expressed in the fetal brain [11], and our earlier biochemical characterization revealed that this protein belonged to the large family of metallophosphoesterases [9, 12]. We could identify orthologs in all mammals and other vertebrates, as well as *Caenorhabditis elegans* and *Drosophila melanogaster* [9, 12]. Indeed, a closely related structural ortholog is found in *Mycobacterium tuberculosis* [13]. There are two variants of the MPPED family in higher organisms, MPPED1 and MPPED2 (239AB) [14], and these two mammalian orthologs show more than 80% similarity at the amino acid level and share similar biochemical properties [9].

The metallophosphoesterases represent a large and diverse group of proteins with similar structural folds that harbor two essential metal ions at the catalytic site [12]. MPPED1 and MPPED2 hydroylze phosphodiester bonds [9]. The substrates for many of these enzymes remain unknown, and while MPPED1 and MPPED2 can hydrolyze 3’,5’-cAMP, the product of this reaction is 3’AMP [9, 15], in contrast to well characterized mammalian cyclic nucleotide phosphodiesterases that produce 5’ AMP as the hydrolysis product [16]. Interestingly, 2’3’-cAMP is the preferred substrate for this group of enzymes, forming 3’ AMP and 2’ AMP as products [17], but the biological relevance of this reaction is unknown. MPPED1 and MPPED2 are also able to hydrolyze a number of colorigenic substrates and such assays revealed that these enzymes could not hydrolyze monoesters but only diester-containing molecules such as bis-p-nitrophenyl phosphate (bis-pNPP) p-nitrophenyl phenylphosphonate (p-NPP) and TMP p-nitrophenyl ester (TMPP) [9].

*D. melanogaster* harbors a single ortholog of MPPED1/MPPED2, which we here call dMPPED [9]. In FlyBase, this gene is annotated as *Fbgn0036028* or *CG16717*. There is no information available on the role of this gene product till date. With a view to decipher the role of *dMPPED* in the fly, we have biochemically characterized this protein, and find that it is a phosphodiesterase. The major sites of expression of *dMPPED* are in the adult brain, testis and ovaries. Mutant flies harboring a deletion of this gene had dramatically reduced lifespan, with no change in fecundity, in an apparently insulin signaling-independent manner. RNA sequencing (RNA-seq) analysis identified a number of pathways which were mis-regulated. Importantly, lifespan could be restored in the mutant fly by neuronal expression of the mammalian MPPED2 ortholog, and neuronal overexpression of dMPPED in wild type flies was sufficient to increase lifespan by ~ 20%. Our findings thus identify a novel gene associated with longevity in the fly that affects a number of signaling pathways associated with aging in the fly.

## Results

### dMPPED is a metallophosphodiesterase expressed during development and in multiple adult tissues

*CG16717* is located on chromosome 3 at cytogenetic location 67C4. The *CG16717* gene has two exons that are spliced to yield a single transcript, encoding for a protein of 300 amino acids. The entire protein coding sequence is present in exon 2 (http://flybase.org/reports/FBgn0036028.html). Since *CG16717* is the only ortholog of the MPPED1/MPPED2 family of proteins in *Drosophila*, we will henceforth refer to it as *dMPPED* (*Drosophila* **m**etallo**p**hos**p**ho**e**sterase **d**omain containing). Sequence alignment of dMPPED, hMPPED1 (human MPPED1), hMPPED2 (human MPPED2) and rMPPED2 (rat MPPED2) reveals that dMPPED shares ~50% sequence similarity to the mammalian orthologs. All the critical residues required to classify dMPPED as a metallophosphoesterase are conserved including the metal binding residue aspartate at amino acid position 49 (D49) and histidine at position 51 (H51; H67 in MPPED2) [15] (Figure 1A).

**Figure 1.**
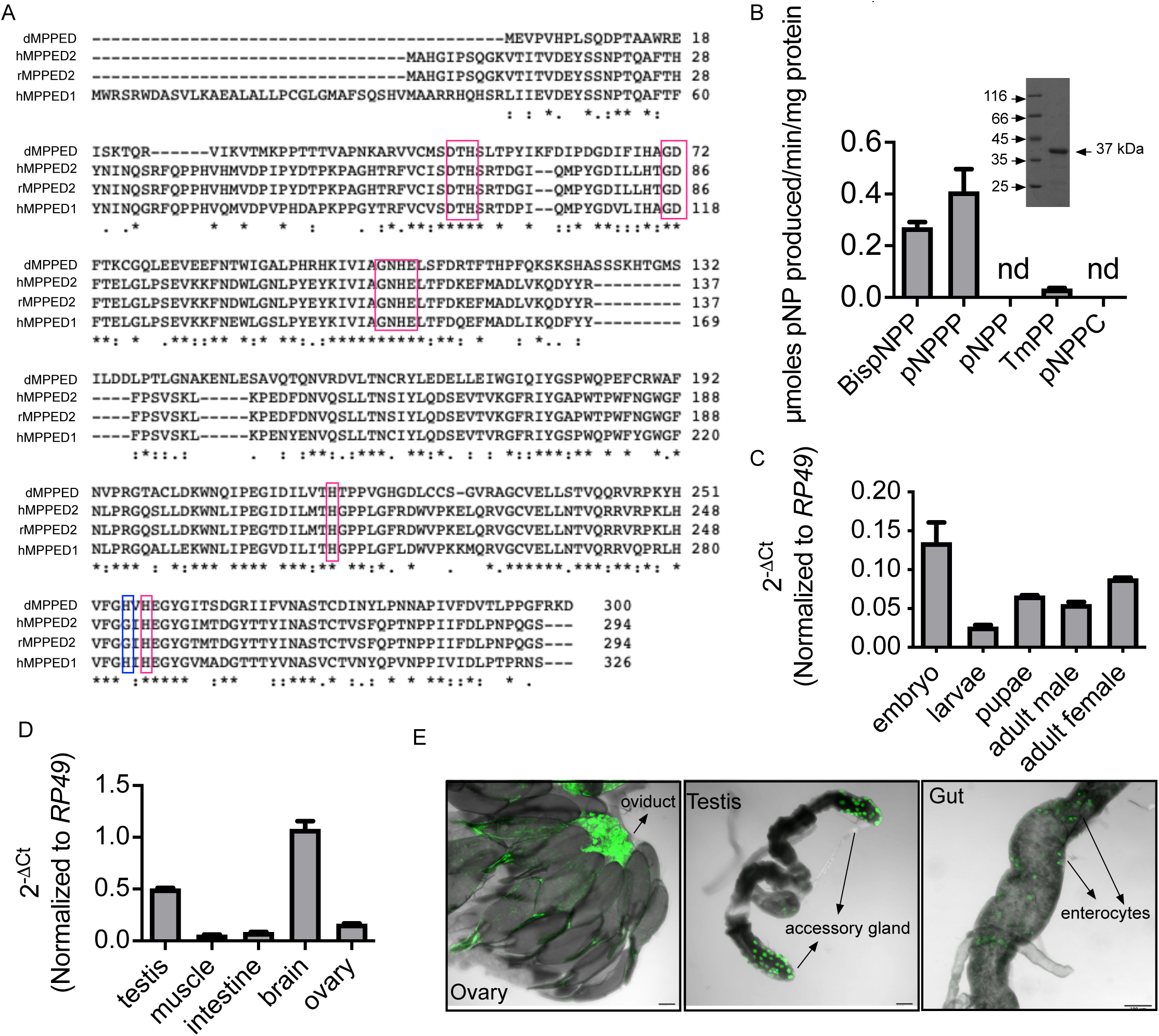
dMPPED is a metallophosphoesterase enriched in the fly neurons. (A) Sequence alignment of dMPPED with human MPPED1, human MPPED2 and rat MPPED2. Highlighted in the blue box is the histidine residue that distinguishes MPPED1 from MPPED2. (B) Catalytic activity of dMPPED with the indicated colorigenic substrates (10 mM), in the presence of Mn^2+^ (5 mM) with dMPPED (500 ng protein). Values represent the mean ± S.E. of duplicate determinations of experiments preformed using two independent protein preparations. BispNPP, bis(p-nitrophenyl) phosphate; pNPPP, p-nitrophenyl phenylphosphonate; pNPP, p-nitrophenyl phosphate; TmPP, thymidine 5’-monophosphate-p-nitrophenyl ester; pNPPC, p-nitrophenylphosphoryl-choline. Inset: a Coomassie R stained gel showing the purified protein used for assays. (C) Expression pattern of *dMPPED* across developmental stages of the fly. Embryos (50 μl; 2 h after egg laying), third instar larvae (10 wandering), pupae (10; 21 h after pupae formation), male and female flies (3 days old); 10 each) were collected and RT-qPCR was performed. *dMPPED* transcript levels have been normalized to *RP49* transcript levels. Graph represents mean ± S.D. from 2 sets of samples collected independently. (D) Expression pattern of *dMPPED* in different adult tissues. Testis and ovaries were collected from 50 male and female flies respectively. Intestine (gut) was dissected from ~ 50 flies in total, and 100 flies were used for muscles and brains. Graph represents mean ± S.D. from 2 sets of samples collected independently. (E) Confocal images of the ovary, testis and posterior midgut (R3-R4) in *pBAC(IT.GAL4)CG16717/UAS-mCD8GFP*. *pBac(IT.GAL4)* enhancer trap element was within the *CG16717* gene. Ovaries show expression in the oviduct, the testis in secondary cells and scattered enterocytes are GFP positive in the posterior midgut. The scale bar indicates 100 μm.

We expressed and purified dMPPED to determine its biochemical properties. The purified protein migrated as a protein of 37 kDa corresponding to the predicted molecular weight (Figure 1B inset). Purified dMPPED was tested for phosphoesterase activity against a panel of colorigenic substrates, and in a manner similar to its mammalian orthologs rMPPED1 and rMPPED2, dMPPED, in the presence of Mn^2+^ as the metal cofactor, showed phosphodiesterase activity and no detectable phosphomonoesterase activity (Figure 1B). dMPPED hydrolysed TmPP poorly, but not pNPPC, indicating some degree of specificity towards the kind of phosphodiester substrates it utilizes which is similar to rMPPED1 and rMPPED2 [9]. A number of divalent cations could be used by the enzyme, in contrast to MPPED2 which showed little detectable activity with Ni^2+^ (Supplemental Figure 1A). The *Km* for Mn^2+^ and pNPPP was 1.8 mM and ~10 mM respectively, comparable to that of MPPED2 [9] (Supplemental Figure 1B, C). dMPPED was active against cyclic nucleotides and hydrolyzed 2’3’cAMP to yield 3’AMP as the product (Supplemental Figure 1D). Unlike MPPED2 which bound 5’GMP with high affinity and was inhibited by it with an IC50 of ~70 nM [15], dMPPED had an IC50 in the range of 400 mM (Supplemental Figure 1D).

To explore the expression profile of *dMPPED* in various stages of development, we performed RT-qPCR. We observed that dMPPED is expressed through all stages of development from embryos to adult flies (Figure 1C). Expression was seen in the adult brain, testis and ovary (Figure 1D), in accordance with expression data gleaned from FlyAtlas2 (Supplemental Figure 2A; http://flyatlas.gla.ac.uk/FlyAtlas2/index.html). We also attempted to utilize genetic approaches to monitor expression, using *IT-gal4^1111-G4^* flies [18]. These flies have the *GAL4* coding sequence inserted within the only intron of *dMPPED* and transcription is therefore expected to be under the control of the same genetic elements as that of *dMPPED*. We crossed these flies to *UAS-mCD8-GFP* flies, adult tissues were dissected and imaged. In agreement with the RT-qPCR data, GFP expression was detected in the adult ovaries in the oviduct, accessory glands of the testis, and to low levels in enterocytes in the intestine (Figure 1E). Interestingly, expression was poorly detected in the brain, perhaps indicating that a neuronal specific enhancer may be present in the intron where the *GAL4* sequence was inserted.

### dMPPED regulates fly lifespan in an IIS/TOR signaling independent manner

To determine the functional role of *dMPPED* in *Drosophila,* we generated a null allele (*dMPPED*^*KO*^) using ends-out homologous recombination [19-21] (Supplemental Figure 3A). The gene deletion was verified by genomic PCR and RT-PCR (Supplemental Figure 3B). In addition, we generated an antibody to dMPPED, and western blotting confirmed the absence of and protein in the heads of *dMPPED*^*KO*^ flies (Supplemental Figure 3C and 3D). To rule out any off-site target effects on adjacent genes, we monitored the expression levels of *α-tubulin* and *furry* and observed that they were not affected in *dMPPED*^*KO*^ flies (Supplemental Figure 3E).

*dMPPED*^*KO*^ flies were homozygous viable and showed no obvious phenotypic differences when compared to wild type flies. However, we noticed that the lifespans of both male and female *dMPPED*^*KO*^ flies were much reduced. We continued to perform lifespan analysis in female flies. Systematic evaluation in female flies demonstrated that the median and maximum lifespan of *dMPPED*^*KO*^ flies were 18% and 20% lower compared to wild type flies (Figure 2A; Supplemental Table 1). We expressed dMPPED ubiquitously using the *Act-GAL4* driver in *dMPPED*^*KO*^ flies, and expression of the transgene was confirmed by western blot analysis (Figure 2A left panel). These flies showed a 41% increase in median lifespan and 47% increase in maximum lifespan over *GAL4* control lines in the mutant background (Figure 2A, right panel). This confirmed that the reduced lifespan exhibited by *dMPPED*^*KO*^ flies was due to loss of dMPPED.

**Figure 2.**
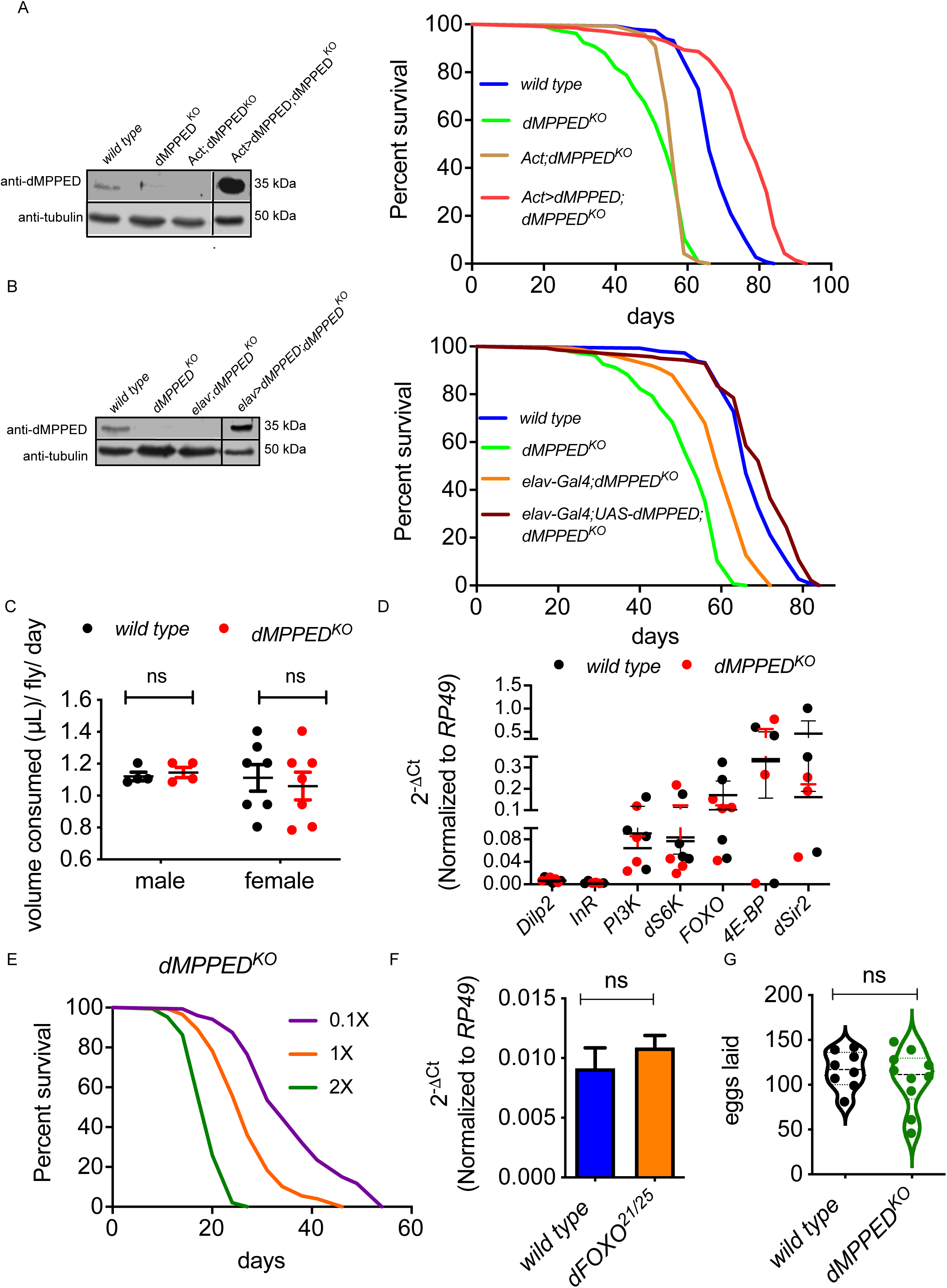
dMPPED regulates fly lifespan in an IIS/TOR signaling independent manner. (A) Left panel: western blot to confirm that *dMPPED*^*KO*^ is a protein null allele, and expression of dMPPED in the rescue line is higher than in wild type flies. The blot was re-probed with anti-tubulin antibody to normalize for protein loading. Right panel: Survival curves of flies of indicated genotypes. (B) Left panel: western blot to detect expression of dMPPED in in indicated lines. The blot was re-probed with anti-tubulin antibody to normalize for protein loading. Right panel: survival curves of indicated lines indicate that neuronal expression of dMPPED rescues lifespan of *dMPPED*^*KO*^ flies. (C) Feeding behaviour of flies measured using the capillary feeder assay (CAFE). Each dot represents a cohort of 5 flies in a capillary feeding chamber. Graph represents mean ± S.D. from two independent experiments. Statistical significance was calculated on GraphPad Prism 5 using the Student’s t-test. (D) Expression levels of some genes in the IIS/ TOR signaling pathway, measured in female wild type and *dMPPED*^*KO*^ flies (10 each). Graph represents mean ± S.D. from 2 independent experiments. (E) Effect of dietary restriction on the survival of *dMPPED*^*KO*^ flies. Flies were subjected to varying concentrations of dietary yeast and their survival curves were plotted. (F) Expression levels of dMPPED in wild type and *dFOXO^21/25^* flies. Transcript levels of *dMPPED* have been normalized to *RP49* transcript levels. Graph represents mean ± S.D. from 2 independent experiments performed using 10 flies in each experiment. Statistical significance was calculated on GraphPad Prism using the Student’s t-test. (G) Fecundity of female wild type and *dMPPED*^*KO*^ flies. The number of eggs laid by flies over a period of 3 days are indicated. Each dot represents data from a single fly. Statistical significance was calculated on GraphPad Prism using the Student’s t-test.

Given that *dMPPED* is expressed at high levels in the brain, and neuronal regulation of lifespan is well studied [22-24], we expressed *dMPPED* using a pan-neuronal driver, *elav-GAL4* (Figure 2B left panel). Interestingly, reconstitution of dMPPED in neurons was sufficient to rescue the reduced lifespan of *dMPPED*^*KO*^ flies (Figure 2B, right panel).

A number of studies have established that dietary restriction (DR) can lead to lifespan extension in multiple model organisms including *Drosophila* [25] and that both very low or very high intake of calories can result in decreased longevity in flies [26]. The reduced lifespan of *dMPPED*^*KO*^ flies could therefore be a consequence of either extreme starvation or increased calorie uptake. The amount of food consumption between wild type and *dMPPED*^*KO*^ flies was tested using the capillary feeder (CAFE) assay [27]. It was found that the volume of food taken up by wild type flies (both male and female) and *dMPPED*^*KO*^ flies was similar (Figure 2C), thus ruling out starvation or increased calorie uptake as the cause for lifespan reduction.

The insulin/insulin-like growth factor (IGF)-1 signaling (IIS) and target of rapamycin (TOR) pathways have been shown to regulate lifespan in *C. elegans, Drosophila*, mice and humans [28]. Binding of the *Drosophila* insulin-like peptide (dILP) to the insulin receptor (InR) leads to phosphorylation and nuclear exclusion of the transcription factor dFOXO via a cascade of signaling events including phosphorylation of the lipid PIP2 to PIP3 by PI3K [29]. Conversely, under conditions of reduced IIS, dFOXO translocates from the cytosol to the nucleus and upregulates target genes involved in longevity [30]. Two such dFOXO target genes that are well studied are S6K and 4EBP. dILP2 is the closest homolog of human insulin [31], has been implicated in regulating fly lifespan and expression is regulated by IIS by a feedback mechanism. In order to test if the reduced lifespan in *dMPPED*^*KO*^ flies was due to mis-regulation of a few components of IIS/TOR pathway, RT-qPCR was performed on whole fly RNA. There was no difference in the transcript levels of *dILP2, InR, PI3K, dFOXO* or the targets *S6K* or *4EBP* in *dMPPED*^*KO*^ flies compared to wild type flies (Figure 2D). However, since many of these proteins are post-translationally modified to increase their activity [28], we took additional approaches to determine if dMPPED regulated lifespan via the IIS/TOR signaling pathway.

Modulating signaling via the IIS/TOR pathway is known to block the increase in lifespan seen upon dietary restriction [25]. We subjected *dMPPED*^*KO*^ flies to food with decreasing yeast concentrations and observed that *dMPPED*^*KO*^ flies showed an increase in median survival. This implied that the IIS/TOR pathway was largely intact in these flies (Figure 2E). Moreover, *dMPPED* was not directly under the control of dFOXO since *dMPPED* levels were similar in wild type and a *dFOXO* hypomorph fly line (Figure 2F). Finally, an increase in IIS/TOR activity often results in an increase in female fecundity [32]. However, the number of eggs laid by *dMPPED*^*KO*^ flies were similar to that laid by wild type flies (Figure 2G). Overall, these results suggest that the reduced lifespan of *dMPPED*^*KO*^ flies was not a consequence of altered IIS/TOR signaling.

### Catalytic activity of dMPPED in not required to regulate lifespan

Metallophosphoesterases can have a wide range of substrates and some even have roles independent of their catalytic activity [12]. We therefore attempted to rescue the reduced lifespan of *dMPPED*^*KO*^ flies by expressing a catalytically inactive version of dMPPED under the control of *elav-GAL4*. The aspartic acid in the first block of residues conserved across all metallophosphoesterases (D49 in dMPPED) is important for metal binding at the active site (Figure 1A) [12]. In mammalian MPPED2, changing the equivalent aspartic acid to an alanine (MPPED2D65A) rendered the protein inactive [9]. Recombinant dMPPEDD49A was expressed and purified, and such a mutation indeed rendered dMPPED inactive (Figure 3A). Transgenic flies expressing the mutant protein were generated and used to test whether they could rescue the reduced lifespan of *dMPPED*^*KO*^ flies. dMPPEDD49A was expressed at lower levels than wild type dMPPED (Figure 3B), suggesting that dMPPEDD49A was intrinsically less stable than the wild type protein, since transcript levels were similar (data not shown). Nevertheless, *dMPPEDD49A* expressed under *elav-GAL4* was able to rescue the reduced lifespan of *dMPPED*^*KO*^flies (Figure 3C; Supplemental Table 1), indicating that dMPPED’s role in regulating lifespan was independent of its catalytic activity. This therefore suggests that dMPPED could be involved in critical interactions with proteins or nucleic acids, as is seen for other members of the Class III metallophosphoesterases [12, 33, 34].

**Figure 3.**
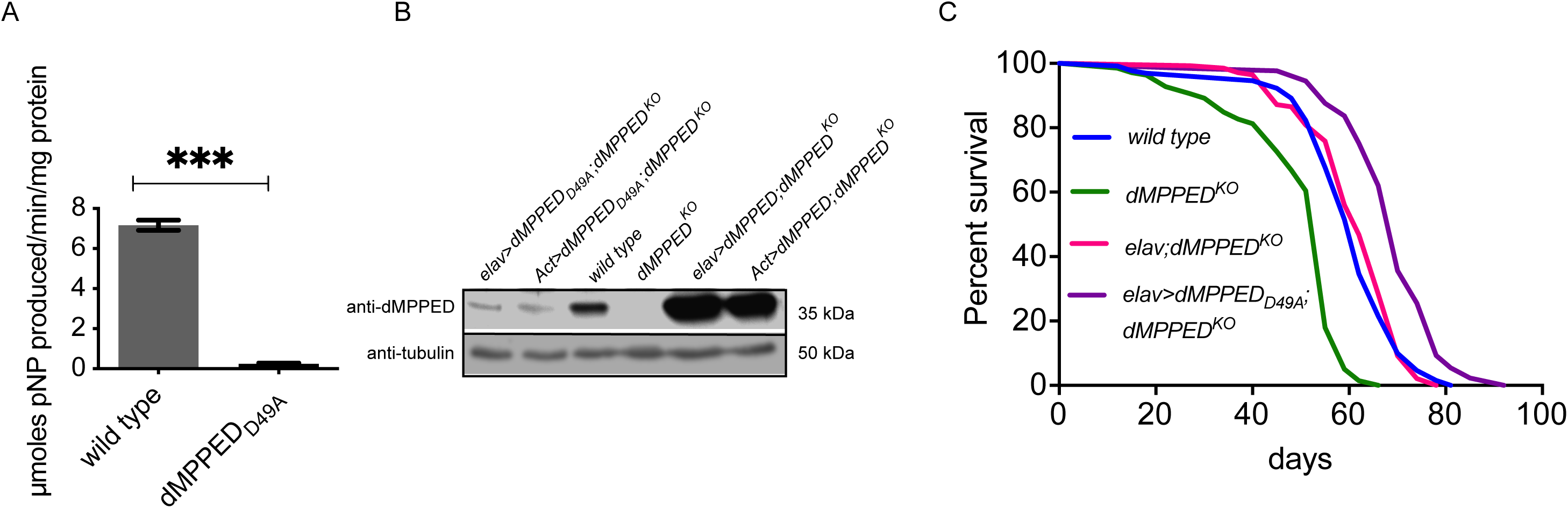
dMPPED regulates fly lifespan in a catalytic activity independent manner. (A) Catalytic activity of wild type dMPPED and dMPPEDD49A. Reactions were performed with 10 mM of p-nitrophenyl phenylphosphonate (pNPPP) in the presence of 5 mM Mn^2+^ and 500 ng of protein. Values represent the mean ± S.D. of duplicate determinations of experiments performed using two independent protein preparations. Statistical significance was calculated on GraphPad Prism using the Student’s t-test. (B) Western blot to detect the expression levels of dMPPED_D49A_. Expression levels of using two different *GAL4* drivers (*Act-GAL4* and *elav-GAL4*) have been shown. The blot was re-probed with anti-tubulin antibody. (C) Survival curves of indicated lines.

Although dMPPEDD49A was able to rescue the reduced lifespan of *dMPPED*^*KO*^ flies, the change in median and maximum lifespan between *elav*>d*MPPEDD49A;dMPPED^KO^* and the corresponding control, *elav;dMPPED^KO^* was 12.9% and 6.5%, respectively. This was lower than that seen with *elav*>*dMPPED;dMPPED^KO^* and the *elav;dMPPED^KO^* control flies which showed a change in median and maximum lifespan of 22% and 13.8%. It is possible that this difference could be a consequence of lower expression levels of dMPPEDD49A, since the efficient functioning of interacting proteins would depend on their concentrations within a cell.

### dMPPED extends the lifespan of wild type flies

Expression of either the wild type or the catalytically inactive versions of dMPPED, rescued lifespan in *dMPPED*^*KO*^ flies, and in fact, marginally extended lifespan (Figures 2, 3). We therefore asked if increasing dMPPED levels in wild type flies could extend lifespan, thereby indicating the sufficiency of *dMPPED* in contributing to lifespan in flies. Over expression of dMPPED using *Act-GAL4* in flies was confirmed by western blot analysis (Figure 4A, left panel) and resulted in an increase in median and maximum lifespan of 22.5% and 23.6% respectively, compared to *Act-GAL4* control flies (Figure 4A, right panel). This extension in lifespan was also achieved upon pan-neuronal expression of *dMPPED* using *elav-GAL4* (Figure 4B). In this case the change in median and maximum lifespan was 17.3% and 14.8% respectively compared to *elav-GAL4* control flies, suggesting that expression of dMPPED in cells other than neurons also contributed to overall lifespan extension in the fly. Over expressing dMPPEDD49A in neurons was sufficient to extend median lifespan by 10.3% and maximum lifespan by 6.9% compared to *elav-GAL4* controls (Figure 4C). This is lower than that seen on over expression of wild type protein, but nonetheless significant. This could again be a consequence of poorer expression of the mutant protein in flies, or that possibility that catalytic activity of dMPPED also plays a role in extension of life span.

**Figure 4.**
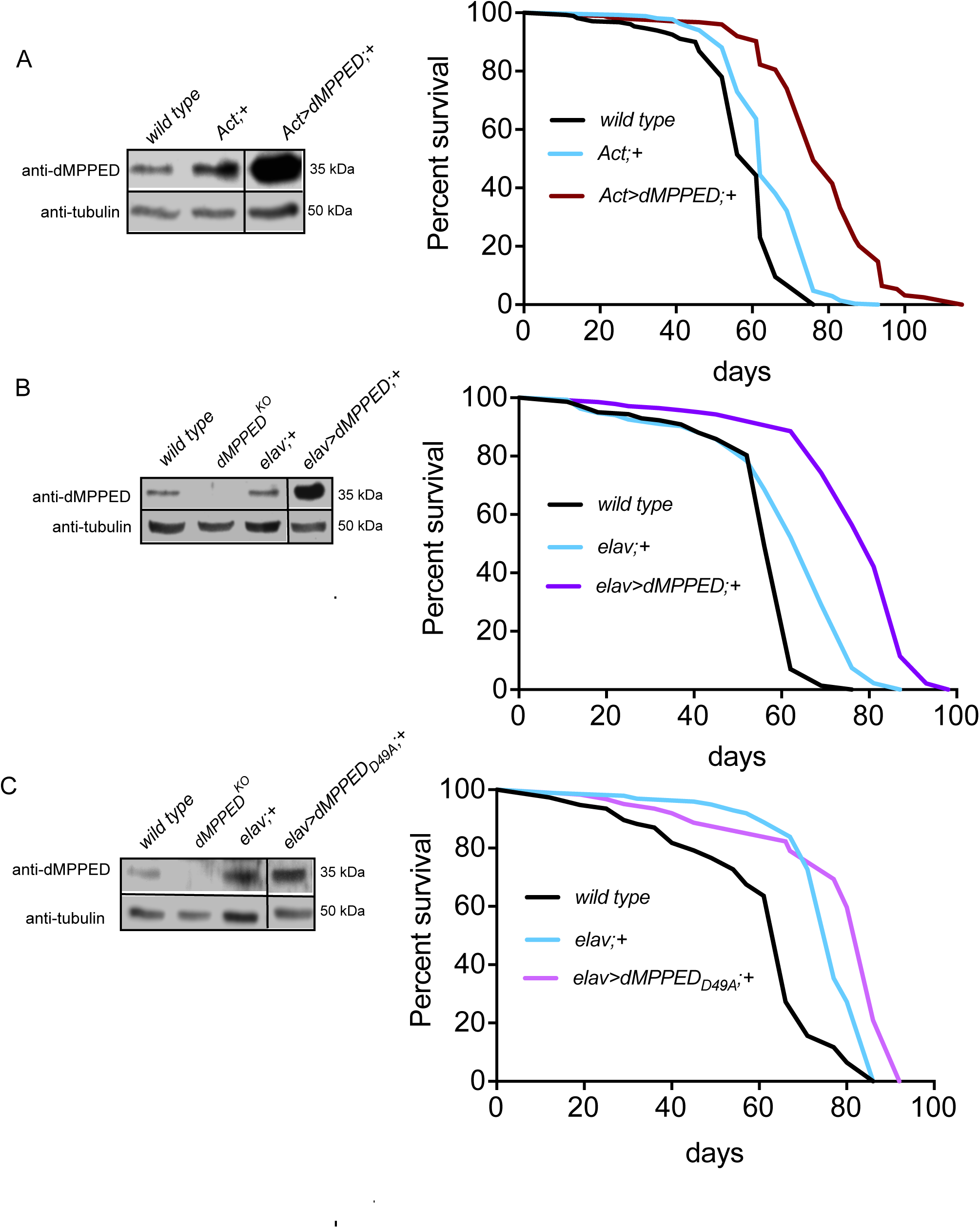
Overexpression of dMPPED extends fly lifespan in a catalytic activity independent manner. (A-C) Left panel: western blot showing the level of expression of dMPPED in different fly lines. The blot was re-probed with anti-tubulin antibody. Right panel: Survival curves indicated fly lines.

### Deficiency of dMPPED mis-regulates genes involved in immunity and regulation of RNA polymerase II activity

We have so far shown that dMPPED is critical for maintaining lifespan in an IIS/TOR-independent manner. Since lifespan is determined by a number of factors that regulate diverse physiological processes, we adopted an unbiased approach to identify genes whose mis-regulation could account for the shortened lifespan in *dMPPED*^*KO*^ flies. We performed RNA-seq analysis of whole flies at 35 days of age, a time where *dMPPED*^*KO*^ flies had started to succumb, but wild type flies showed >99% survival. A number of genes were differentially regulated in *dMPPED*^*KO*^ flies (Figure 5A), with the majority of them being of unknown function. A number of mis-regulated genes are co-expressed with dMPPED in different tissues (Figure 5B and Figure 1D).

**Figure 5.**
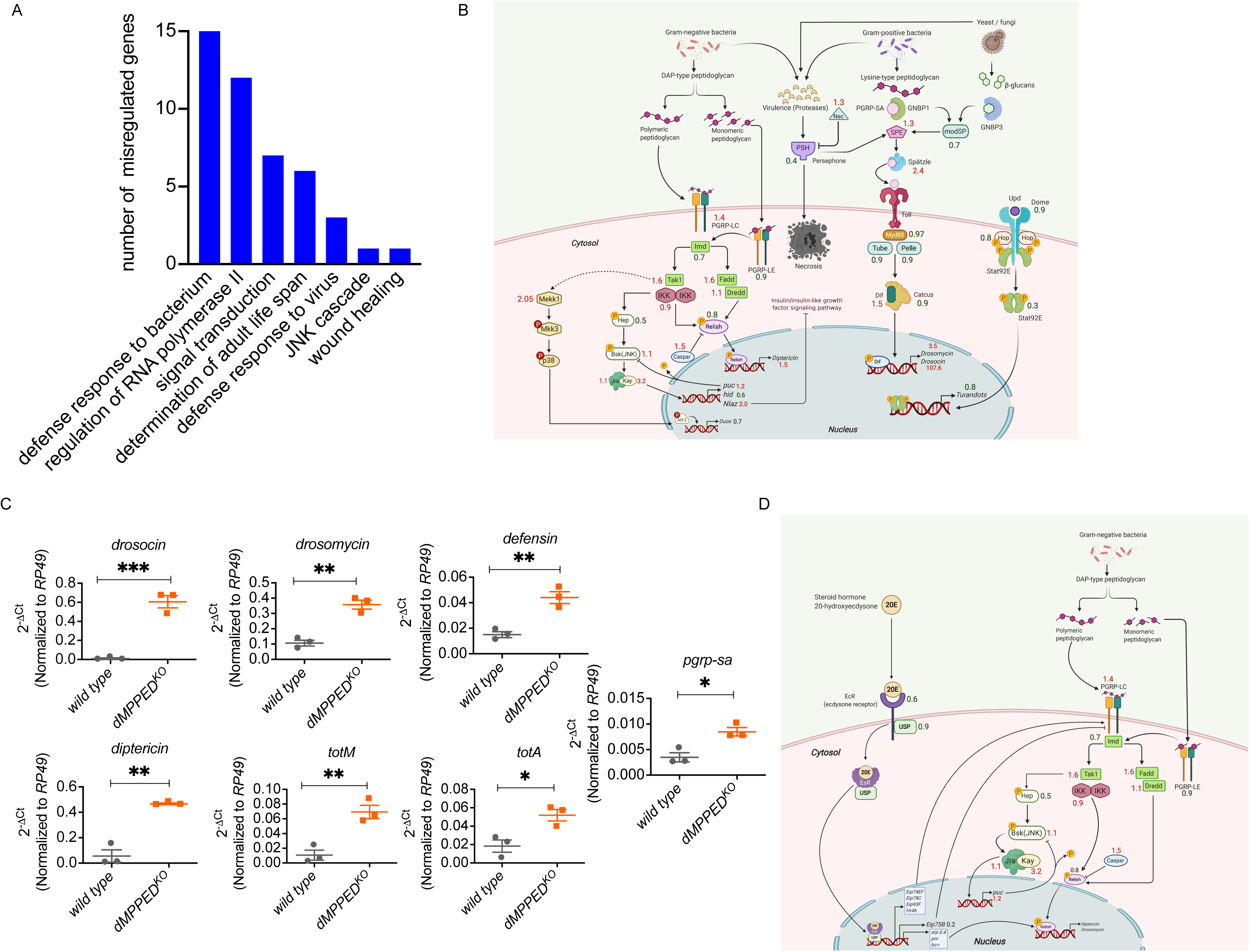
RNA-seq analysis and mis-regulation of IIS pathway genes. (A) A volcano plot showing protein-coding genes altered in 35-day old *dMPPED*^*KO*^flies. The Log_2_ fold change (Log_2_FC) values are plotted on the x-axis and negative log (base 10) of the P value (-Log_10_P) on the y-axis. Blue dots represent genes with P-values <0.05 and Log2 fold change values of more or less than one fold. (B) The tissue distribution of differentially regulated genes. Misregulated genes (852, with 592 up-regulated and 260 down-regulated) were analysed by DGET (http://www.flyrnai.org/tools/dget/web/) based on published analysis of the Drosophila transcriptome [92]. (C) The IIS/TOR pathways in *Drosophila*. Numbers indicate the fold-change in genes from RNA-seq analysis with numbers in red indicating upregulated genes, and those in grey downregulated genes. The figure has been created with Biorender.

Interestingly, a few genes that have been shown to regulate lifespan showed a pattern of expression in *dMPPED*^*KO*^ flies that correlated with earlier findings. For example, overexpression of *sun* (*stunted,* a ligand of *methuselah;* fold change 2.7; [35]) and *puc* (important in JNK signaling; fold change 1.2; [36]) would result in shortened lifespan, based on earlier reports. Similarly, down regulation of *magu [37], Gnmt [38], cct1* [39]*, l(3)neo18* [40] and *rb* [41] in independent studies are associated with a decrease in lifespan; all these genes were down regulated in *dMPPED*^*KO*^ flies (fold change from 0.2 to 0.8).

RNA-seq analysis also revealed that genes in the IIS pathway were mis-regulated in 35 day-old *dMPPED*^*KO*^ flies (Figure 5B). For example, *Inr* showed a 40% reduction in transcript levels in *dMPPED*^*KO*^ flies, and *chico* and *PI3K* were significantly upregulated. Such changes in gene expression were not detected in day 3 flies (Figure 2D) and are not reported in aging flies [42]. Neural Lazarillo (*NLaz*) is an extracellular lipid binding protein of the lipocalin family and its expression is regulated by the JNK signaling pathway [43]. Overexpression of *NLaz* extends lifespan by repressing IIS activity in adult flies. In *dMPPED*^*KO*^ flies, *Nlaz* showed 2-fold higher expression than wild type flies, suggesting that may serve to reduce IIS signaling. Both the downregulation of *InR* and upregulation of *Nlaz* may have offset the effects of increased *chico* and *PI3K* expression in *dMPPED*^*KO*^ flies, but it is interesting to note that the IIS pathway seems to be perturbed in aging, but not younger, flies deficient in dMPPED. We then sought to categorize genes of known function in pathways using Gene Ontology with significant fold regulation (>1.2 and < -0.8) (Figure 6A). A large set of genes important in defense response to bacterium were also mis-regulated in *dMPPED*^*KO*^ flies, as were a few genes important for viral resistance and wound healing (Figure 6B). Changes in levels of genes in the IMD and Toll pathways could account for the upregulation of AMP expression in *dMPPED*^*KO*^ flies. These genes included those for the antimicrobial peptides (AMPs) such as *diptericin, drosomycin* and *drosocin*. We validated the RNA-seq analysis by measuring transcript levels by RT-qPCR and found that a number of genes associated with innate immune pathways in flies are induced (Figure 6C).

**Figure 6.**
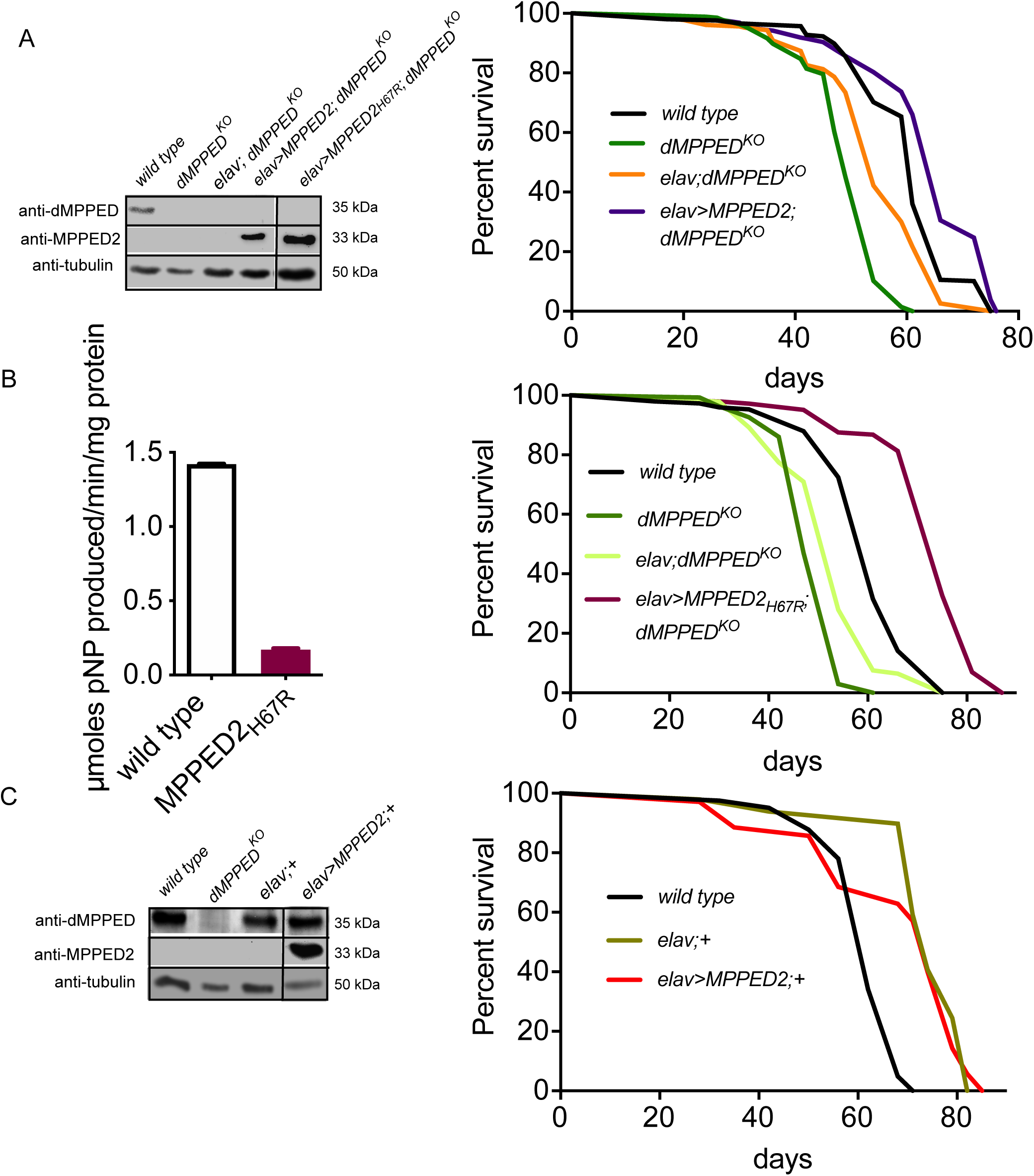
Mis-regulation of genes involved in innate immunity in *dMPPED*^*KO*^ flies. (A) Differentially regulated genes with P value cut off < 0.05 and either 1.2 or -0.8 Log_2_FC were analysed in GO functions-biological processes (http://geneontology.org). Care was taken to ensure that genes were not represented more than once in each pathway. (B) Innate immunity pathways in *Drosophila* indicating genes whose expression was significantly altered based on RNA-seq analysis. Numbers indicate the fold-change in genes from RNA-seq analysis with numbers in red indicating upregulated genes, and those in grey downregulated genes. (C) RT-qPCR to confirm upregulation of AMPs and other genes involved in innate immunity in *dMPPED*^*KO*^ flies. (D) Pathway delineating the process by which 20E regulates AMP production via PGRP-LC. Numbers indicate the fold-change in genes from RNA-seq analysis with numbers in red indicating upregulated genes, and those in grey downregulated genes. The figures have been created with Biorender.

PGRP-LC, the peptidoglycan-recognition protein LC encodes a receptor that recognizes diaminopimelic acid-type peptidoglycan and is important for the Imd-mediated induction of AMPs [44]. RNA-seq analysis reveals that it is upregulated 1.4-fold in *dMPPED*^*KO*^ flies. *PGRP-LC* is upregulated by 20-hydroxyecdysone (20-E) in adult flies [45], a hormone central to development and metamorphosis in the fly (Figure 6D). 20-E binding to its receptor increases the transcription of a number of genes including *Eip75B* and *kayak (kay)* both of which regulate AMP production [45]. We find that *kay* levels are increased 3.2-fold in *dMPPED*^*KO*^ flies. Moreover, *Eip75B* [46], which is a negative regulator of PGRP-LC [45, 47], is down-regulated significantly in *dMPPED*^*KO*^ flies. The mis-regulation of both these genes could account for the increase in AMP transcripts seen in *dMPPED*^*KO*^ flies.

This early life inflammation in the absence of infection is a hallmark of faster aging and shortening of lifespan [48-50] and could contribute to the reduction in lifespan seen in *dMPPED*^*KO*^ flies. Interestingly, *dMPPED* has been shown to be one of the major genes upregulated in S2 cells following infection with *Drosophila* C virus [51]. Infection with this virus activated the JAK/STAT and Imd pathways, and it is interesting to note that a few genes in these pathways are misregulated in *dMPPED*^*KO*^ flies (Figure 6A). Thus, these findings appear to place dMPPED as a link between two important innate immune pathways in the fly.

A number of genes are regulators of RNA polymerase II activity and therefore serve as DNA-binding transcription factors. Of interest is *HmgZ*, upregulated 7-fold, is expressed in the nervous system of the larvae and in adults and genetically interacts with the Brahma chromatin complex [52]; *gce* (germ cell expressed) is a basic helix-loop-helix-PAS domain containing transcription factor and is a paralog of *Met* [53]. *Gce* encodes the receptor for sesquiterpenoid juvenile hormone [54] and is upregulated ~ 3-fold in *dMPPED*^*KO*^ flies. Interestingly, it has been recently shown that Gce and Met signal to intestinal progenitors to generate a larger organ to assist in increased lipid metabolism required for reproductive output [55]. *maf-S* (up-regulated 2.5-fold) heterodimerizes with Nrf2 (not changed in our data set), and its overexpression has been shown to preserve Nrf2 signaling to antagonize age-associated functional decline [56], an observation which appears to be contradictory to our findings.

Dorsal (*dl*) is a member of the NF-kappaB/Rel family of proteins and its 2-fold higher levels in *dMPPED*^*KO*^ flies could account for the increased expression of *drosomycin* and *defensin* [57]. Finally, *tou*, upregulated 2-fold, activates neural gene expression via interaction with *pnr* (not significantly changed in our data set) and has roles in nervous system and lymph gland development [58].

In summary, RNA-seq uncovered the potential of *dMPPED2* to regulate a number of pathways that could affect lifespan in the fly, ranging from immune genes to core transcription factors. Moreover, many of these genes are expressed in tissues where dMPPED is detected, suggesting multiple tissue-specific changes in the knock-out flies that could act in concert to regulate lifespan.

### Mammalian ortholog of dMPPED regulates lifespan in flies in an activity independent manner

The MPPEDs are highly evolutionarily conserved proteins across metazoans and the sequence similarity between the fly and the mammalian MPPED proteins is ~50% (Figure 1A). The reduced lifespan of *dMPPED*^*KO*^ flies offered a good model to test possible functional conservation between dMPPED and its mammalian ortholog, MPPED2. To this end, we expressed rat MPPED2 neuronally in *dMPPED*^*KO*^ flies and confirmed expression by western blot analysis using an MPPED2 specific monoclonal antibody (Figure 7A, left panel). Interestingly, lifespan in flies expressing MPPED2 was restored to wild type levels and resulted in a 22.2% extension in median lifespan and a 13.6% extension in maximum lifespan compared to *elav;dMPPED^KO^* control flies (Figure 7B, right panel). The inactive form of the mammalian protein, MPPED2H67R, which arises from a single nucleotide polymorphism [9], was also able to rescue the lifespan of *dMPPED*^*KO*^ flies (Figure 7B). The *elav*>*MPPED2H67R; dMPPED^KO^* flies showed a median survival of 38.8% and maximum survival of 22.7% greater than the *elav;dMPPED^KO^* control flies (Figure 7C). However, overexpression of MPPED2 in wild type flies failed to extend lifespan (Figure 7D). This suggests that the reduced lifespan of *dMPPED*^*KO*^ flies and the increased lifespan seen in flies over expressing dMPPED may be regulated by two different mechanisms, and that mammalian MPPED2 has only partial functional conservation with dMPPED.

**Figure 7.**
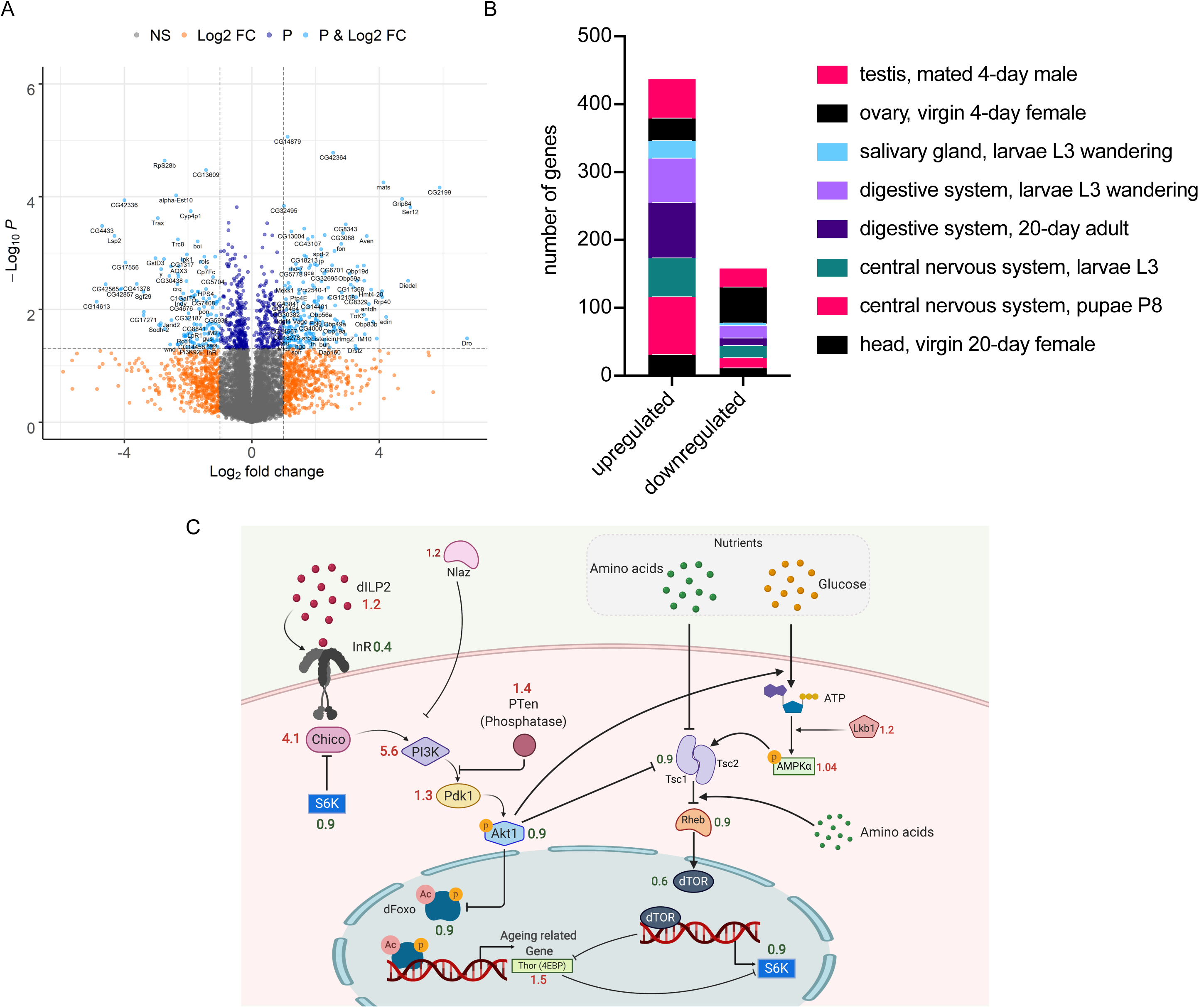
Mammalian ortholog of dMPPED regulates lifespan in flies in an activity independent manner. (A) Left panel: western blot showing expression levels of dMPPED, rat MPPED2 and rat MPPED2_H67R_ in indicated fly lines. The blot was re-probed with anti-tubulin antibody. Right panel: survival curves of indicated fly lines. (B) Left panel: catalytic activity of wild type MPPED2 and dMPPED2_H67R_. Reactions were performed with 10 mM of p-nitrophenyl phenylphosphonate (pNPPP) in the presence of 5 mM Mn^2+^ and 500 ng of protein. Values represent the mean ± S.E. of duplicate determinations of experiments preformed using two independent protein preparations. Right panel: survival curves of indicated fly lines. (C) Left panel: western blot showing expression levels of dMPPED and MPPED2 in the indicated fly lines. The blot was re-probed with anti-tubulin antibody. Right panel: Survival curves of indicated fly lines.

## Discussion

Here, we present the first characterization of the fly ortholog of mammalian MPPED1/MPPED2 proteins and show its importance in maintaining the normal lifespan of the adult fly. While a number of genes that regulate lifespan in the fly are also critical for normal development, it appears that dMPPED is not essential during development, but plays a role in aging in the adult fly. Importantly, the catalytic activity of dMPPED is not essential for its role in modulating lifespan, and it probably functions as a scaffolding protein. dMPPED is able to hydrolyze 2’3’-cAMP to 3’AMP in a manner similar to the mammalian orthologs. 2’3’-cAMP is hydrolyzed by CNPase in mammals and accumulation of this cyclic nucleotide leads to an increased susceptibility to brain injury and neurological disease [59]. Whether an additional role of dMPPED relates to its catalytic activity, and whether 2’3’ cAMP has a role to play in the fly brain remains to be investigated.

From analysis of the *Drosophila* interactome described by high throughput approaches [60], only one gene product, SH3PX1, was reported to interact with dMPPED with some degree of confidence. This protein is the single fly ortholog of the human sorting nexin 9 family that is known to function in vesicular sorting [61, 62]. SH3PX1 has been found to regulate the formation of lamellipodia, tubules and long protrusions in S2 cells [63]. In the fly, SH3PX1 localizes to neuromuscular junctions where it regulates synaptic ultrastructure [64]. Neurotransmitter release was significantly diminished in SH3PX1 mutants and functional interactions with Nwk, a conserved F-BAR protein that attenuates synaptic growth and promotes synaptic function in *Drosophila*, was observed [65]. Interaction with Nwk was via the SH3 domain of SH3PX1, that recognizes proline-rich sequences. The co-expression of dMPPED and SH3PX1 in neuronal tissue may allow for functional interaction that could modulate Nwk action, since the SH3 domain in SH3PX1 could interact with the PXXP motif in dMPPED (residues 4-7; Figure 1A) and sequester it from Nwk.

dMPPED shows a marginally higher sequence identity to hMPPED1 than hMPPED2 (47.6% to 48.7%). Are there proteins that have been shown to interact with MPPED1/MPPED2 which may have implications as interacting partners of dMPPED? In an extensive screen across the human genome involving affinity purification and mass spectrometry of interacting proteins (https://bioplex.hms.harvard.edu), the only common proteins that interacted with both MPPED1 and MPPED2 were transcription factors NR2F1 and NR2F6. These nuclear receptor factors are not conserved in *Drosophila*, which has far fewer nuclear-receptor genes that any other model organism [66]. However, a close homolog of these proteins in mammals is NR2F3, or COUP-TF1, which has the ortholog seven up (*svp*) in the fly [66]. *svp* has crucial roles in neuronal development during embryogenesis and in the development of photoreceptor cells [67]. *COUP-TF1* is also required for neuronal development and axon guidance [68]. Interactions, either direct or genetic, between dMPPED and *svp* would be an interesting line of study in future.

*dMPPED* is a pro-longevity gene, which reduces lifespan upon loss-of-function and extends lifespan when overexpressed. A number of genes have been shown to extend lifespan on overexpression in the fly [69] and include *dTsc1* and *dTsc2* (orthologs of tuberous sclerosis complex genes) [70]; *dPTEN* (phosphatase and tensin homolog) [30]; genes encoding enzymes involved in energy homeostasis such as *d-NAAM* (nicotinamide amidase) [71], *men* (malic enzyme) [72], and *zw* (homolog of G6PDH) [73]. In many cases, neuronal expression of genes also extends lifespan, as is seen in the case of *dMPPED* [69]. Overexpression of proteins important in DNA and protein repair, proteolysis, antioxidant activity, and autophagic processes also extend lifespan and include dSir2, dFOXO [30], hSO*D* [74] and MsrA and MsrB (methionine sulfoxide reductases) [75], and the E3 ubiquitin ligase, parkin [76]. dMPPED appears to be unique in that its enzymatic activity is not required to show effects on longevity in contrast to the examples cited above. In RNA-seq analysis, most of these genes showed similar expression in wild type and *dMPPED*^*KO*^ flies, except for *dSir2*, which was downregulated ~ 70%. The extension of lifespan in *dSir2* overexpressing flies seems to be dependent on the genetic background of the flies [77]. However, suppression of *dSir2* expression in neurons resulted in a reduction of lifespan ~ 10-30% [77]. Therefore, the reduction in *dSir2* may contribute to the reduction in lifespan seen in *dMPPED*^*KO*^ flies. It is important to note that there was no significant change in *dSir2* levels in 3day-old flies, suggesting that the impact of sirtuin activity could be more critical as flies age.

The high levels of AMP production in the absence of any overt infection (Figure 6C) is similar to the sustained increase in innate immune activity that is associated with aging in mammals, known as inflammageing, mediated by NF-κB [78]. Recently, aging associated NF-κB dependent AMP gene expression was seen in the head and the brain, specifically in glial cells, and reducing the activity of NF-κB in the brain prolonged life span [79]. However, an upregulation of *Caspar*, a negative regulator of NF-κB, and a modest decrease in *NF-κB* (*relish*) transcripts in *dMPPED*^*KO*^ flies (Figure 6B), indicates a more complex, and perhaps, tissue-specific regulation of AMP production in these flies.

In summary, we have identified a new player in regulating longevity in flies. The presence of orthologs of dMPPED in higher animals suggests that the role of this protein in mammals is worthy of study. Our analysis reveals that this protein could serve as a focal point for interaction and cross talk with a number of pathways, either through direct interaction or by modulating the activity of a few proteins which could then impact more globally.

## Materials and Methods

### Cloning and mutagenesis of dMPPED and MPPED2

The full-length coding region of *dMPPED* was amplified from cDNA prepared from whole flies using dMPPED_MfeI_Fwd (5’ GTCAATTGATATCAAAATGGAAGTG 3’) and dMPPED_XhoI_Rvs (5’ ATCCTCGAGGCATGCTAATCCTTG 3’) primers, digested with XhoI and cloned into EcoRV and XhoI digested pBKSII vector to generate pBKS-dMPPED. The clone was verified by sequencing (Macrogen, South Korea). The MfeI-XhoI fragment from pBKS-dMPPED was cloned into EcoRI and XhoI digested pPROExHT-B vector to obtain pPRO-dMPPED. The same fragment was cloned into EcoRI and XhoI digested pUAST-attB vector to obtain pUAST-attB-dMPPED.

The D49A mutation was generated on pBKS-dMPPED using the dMPPED_D49A_Fwd primer (Supplemental Table 2) following the single oligonucleotide-based mutagenesis protocol [80] to obtain pBKS-dMPPEDD49A. Using the same cloning strategy as the wild-type, pPRO-dMPPEDD49A and pUAST-attB-dMPPEDD49A were generated.

The pBKS-MPPED2 and pBKS-MPPED2H67R plasmids were available in the laboratory [9]. A BamHI-KpnI fragments from these two clones were cloned into BglII and KpnI digested pUAST-attB independently to obtain pUAST-attB-MPPED2 and pUAST-attB-MPPED2H67R, respectively. The pPRO-MPPED2 and pPRO-MPPED2H67R were as reported earlier [9].

### Expression and purification of dMPPED and MPPED2

The pPRO clones of dMPPED were transformed into *E. coli* BL21DE3 and proteins were expressed as described earlier [9]. Gel filtration of purified protein was carried out in buffer containing 50 mM Tris/HCl, 5 mM 2-mercaptoethanol, 50 mM NaCl, and 10% glycerol at pH 8.8 and 4 °C at a flow rate of 200 ul/min using a Superose 12 column and an AKTA fast protein liquid chromatography system (GE Healthcare). The protein eluates were stored in aliquots at -70 °C until further use.

### Biochemical assays

Enzyme assays for various activities (phosphatase, phosphodiesterase, nuclease and phospholipase) were performed in a triple buffer system (MES, HEPES, diethanolamine, 50 mM (pH 9.0)), 5 mM 2-mercaptoethanol, and 10 mM NaCl in the presence of 10 mM concentrations of the specified substrate and 5mM Mn^2+^ as the metal cofactor. Assays were stopped by the addition of 10 ml of 200 mM NaOH, and absorbance was monitored at 405 nm. The amount of p-nitrophenol formed was estimated based on its molar extinction coefficient of 18,450 M^-1^ cm^-1^.

Hydrolysis of 2’,3’-cAMP was analyzed in the same buffer system in the presence of 1 μg of dMPPED, 10 mM of the 2’,3’-cAMP as substrate and 5mM Mn^2+^ cofactor for 120 min. The reaction was terminated by the addition of 50 ul of 20 mM (NH4)H2PO4 (pH 6.2). Aliquots from assays were applied to a Supelcosil LC-8DB (25 cm x 4.6 mm, 5 um) HPLC column equilibrated with 20 mM (NH4)H2PO4 (pH 6.2) at a flow rate of 1 ml/min on an Agilent Technologies HPLC system. The column was calibrated using 2’,3’-cAMP, 3’-AMP and 2’-AMP nucleotides. Inhibition assays were performed in the presence of varying concentrations of 5′-GMP in the assay mix containing 30 mM p-nitrophenyl phenylphosphonate as substrate and 5mM Mn^2+^ as cofactor.

### Fly culture

Unless mentioned otherwise, flies (*Drosophila melanogaster*) were reared on standard media containing cornmeal, dextrose, yeast and agar along with antibacterial and antifungal agents. Cultures were maintained at 25 °C and 50% relative humidity under 12h light/12h dark cycles. The wild type strain used was *w^1118^*. All flies used in this study are listed in Supplemental Table 3.

For dietary restriction (DR) assay, an established protocol was adapted [81]. Flies were maintained on media consisting of 5% sucrose and 1% agar, supplemented with varying concentrations of Brewer’s yeast (0.1X, 1X and 2X with 1X being 100g/L).

Virgin female flies (0-3 day old) were collected and kept in groups of 10 flies per vial. The number of dead flies was recorded every 3 days, when flies were transferred to fresh media vials, for lifespan analyses.

### Dissection and Imaging

Adult tissues were dissected in cold PBS and fixed in 4% paraformaldehyde (PFA) for 20 minutes at room temperature. Samples were then washed thrice with PBS containing 0.1% Triton X-100 for 5 min each and stained with Hoechst nucleic acid stain for 20 minutes. Samples were mounted on a glass slide using Antifade solution and coverslips were sealed using nail-polish. Images were acquired using a Leica TCS SP8 confocal microscope.

### Generation of *dMPPED*^*KO*^ flies

A loss-of-function mutant was generated using ends-out homologous recombination [82]. For this, 4.5kb genomic region immediately upstream of the *dMPPED* coding sequence and 4.5kb genomic region immediately downstream were amplified using fly genomic DNA as template and ExTaq polymerase (Takara). The two amplicons were sequentially cloned into the 5’ and 3’ multiple cloning sites of the pGX-attP vector [83] respectively, thus obtaining pGX-attP-dMPPED. This construct was microinjected into *w^1118^* embryos to obtain P ‘donor’ flies [82]. These flies were used in a series of crosses and the progeny screened for loss of *dMPPED* as described previously [83]. The *dMPPED* knock-out thus obtained was further verified by genomic and RTCPR.

### Genomic and Real time PCR

For genomic DNA isolation, 10 flies were homogenised in 100 μL of buffer-A (100 mM Tris-HCl (pH 7.5), 100 mM EDTA, 100 mM NaCl, 0.5% SDS). An additional 100 μL of buffer-A was added and the samples were incubated at 65 ^°^C for 30 min. To this, 400 μL of buffer-B (1 part 5 M potassium acetate and 2.5 parts of 6 M lithium chloride) was added and the mix was incubated on ice for 10 min. Following this, the debris was removed by centrifugation and the genomic DNA in the supernatant was precipitated using 300 μL of 2-propanol, washed with 70% ethanol, air dried and resuspended in 20 μL of TE (10 mM Tris-HCl (pH 7.5), 1 mM EDTA) and genomic DNA was quantified by measuring the absorbance at 260 nm on a NanoDrop spectrophotometer (Thermo Scientific). Genomic PCR was performed using 50 ng of genomic DNA, 2.5 pmoles of gene specific forward and reverse primers, 0.2 mM dNTPs and 1 U of Taq DNA polymerase in a 20 μL reaction containing 1X standard Taq buffer.

RNA was isolated from flies of indicated age using the TRI reagent (Sigma). Real time quantitative PCR (RT-qPCR) was performed using the VeriQuest SYBR Green qPCR master mix with ROX (Affymetrix) on an ABI 7000 real time PCR machine (Applied Biosystems). Transcript levels of all the genes tested were normalized to transcript levels of ribosomal protein-49 (RP49) using the ΔCt method wherein Ct stands for Cycle Threshold. Transcript levels have been plotted as 2^-ΔCt^ wherein ΔCt = Ctgene -CtRP49.

### Western blot

Purified dMPPED protein was injected into rabbits to raise polyclonal antibody to the protein. Polyclonal antibody against MPPED2 was available in the laboratory [9]. Fly brains were homogenized in 20 μL of homogenization buffer (50 mM Tris-Cl (pH 7.5), 2 mM EDTA, 1 mM DTT, 100 mM NaCl, and 1X Roche protease inhibitor mix). Samples were centrifuged at 13,000 g for 10 min, Laemmli sample buffer was added to the supernatant and boiled for 5 min. Protein samples were resolved on a 12% SDS-polyacrylamide gel and transferred onto a PVDF membrane (Immobilion-P, Millipore). The PVDF membrane was rinsed with TBST (10 mM Tris-HCl (pH 7.5), 100 mM NaCl, 0.1% Tween 20) and blocked for 1 h using 5% BSA made in TBST. Polyclonal anti-dMPPED IgG (1 μg/mL) or anti-MPPED2 (culture supernatant from hybridoma at 1: 500 dilution) was added into the blocking solution and the blot was incubated overnight at 4 °C. The membrane was washed and then incubated with TBST containing 0.2 % BSA and horse radish peroxidise conjugated anti-rabbit secondary antibody for 1h at room-temperature. Bound antibody was detected by Immobilon Western chemiluminescent HRP substrate (Millipore) on the FluorChem Q MultiImage III system (Alpha Innotech). Anti-tubulin antibody (12G10; Developmental Studies Hybridoma Bank) (1:1000) was used to detect tubulin which served as the loading control for individual lanes.

### Capillary feeder assay

Food ingested by flies was quantified using capillary feeder assay (CAFE) [84]. The upper chamber housed the flies and was fitted with a rubber cork holding the calibrated glass micropipette. Calibrated glass micropipettes of 50 μl volume filled with the liquid medium by capillary action were inserted through the cork. A mineral oil overlay of ≈ 0.5 μl was used to minimize evaporation. The lower chamber was half-filled with water. The upper and the lower chamber were sealed using PVC tape, separated by a layer of muslin cloth to keep the upper chamber hydrated. Experiments were conducted at 25°C under 12-hour light/dark cycle. 2-4 day old virgin flies were used and the amount of media consumed from the micropipette was measured after 24 hours of feeding.

### Egg laying

Single virgin female-single male crosses were set up using 4 day old flies and allowed to mate for a period of 12 h. Female flies were transferred to a fresh vial containing egg laying media (1% sucrose, 1% agar, 0.4% propionic acid and a patch of live yeast paste). These mated female flies were maintained at 25 ^°^C under a 12 h light/dark cycle and transferred every 24 h to a fresh vial containing egg laying media. Number of eggs laid per female per day was counted from each vial.

### RNA-Sequencing

10 virgin wild type and *dMPPED*^*KO*^ female flies (35 days old) were anaesthetized and collected. The collection and RNA extraction was done in triplicates. RNA extraction was performed with the help of RNeasy kit (Qiagen) as per manufacturer’s protocol, DNased, and quantified using Nanodrop. 1.5 µg of RNA was subjected to RNA-Sequencing (Genotypic Technology Pvt. Ltd, Bangalore, India). RNA QC was confirmed by Bioanalyzer. The RNA library was prepared as per NEBNext Ultra directional RNA library prep kit. Illumina HiSeq paired end sequencing was performed.

The quality of RNA-Seq reads in the Fastq files of each sample was checked using the FastQC program (v.0.11.4) [85]. The quality of raw reads was measured using quality scores (Phred scores), GC content, per base N content, sequence length distributions, duplication levels, overrepresented sequences, and K-mer content as parameters. Trimmomatic (v.0.36) was used to remove adaptors, and low-quality sequences to rid the raw reads of any artefacts [86]. After filtering, the paired-end reads from each sample were mapped to the reference genome of *Drosophila melanogaster* (Dmel_Release_6) using HISAT2 program (v.2.0.5) [87, 88]. The index for the reference genome required by HISAT2 to identify the genomic positions of each read was provided by downloading the prebuilt index for *D. melanogaster* from the HISAT2 site (http://ccb.jhu.edu/software/hisat2/manual.shtml).

Transcript assembly and relative abundances of isoforms were determined using StringTie (v.2.1.1) [89]. The merged transcripts were fed back into StringTie to re-estimate the transcript abundances using the merged structures. The read counts from this were normalized against gene length to obtain FPKM (Fragment Per Kilobase of exon model per million mapped reads) values using Ballgown [90] and TPM (Transcript Per Million) values were obtained. All FPK values in a particular sample were added up and divide by 1,000,000 to get a “per million” scaling factor. Each of the FPK values were then divided by the “per million” scaling factor to obtain the Transcript Per Million (TPM) values. Genes and transcripts that were differentially expressed between the two genotypes were determined using DESeq2 (V1.26.0) [91]. The resulting P values were adjusted using the Benjamini and Hochberg’s statistical test to control the false discovery rate (FDR) and the volcano plot was generated.

Some bioinformatic analysis was performed by DeepSeeq Bioinformatics, Bengaluru, India.

## Acknowledgements

We thank Dr. Richa Tyagi and Dr. Upendra Nongthomba for initiating these studies. Microinjection of flies was performed in the Fly Facility at NCBS. Funding from the DBT-IISc Partnership Program Phase-II BT/PR27952/INF/22/212/2018/21.01.2019 is acknowledged. SSV is a JC Bose National Fellow (SB/S2/JCB-18/2013) and a Margdarshi Fellow supported by the Wellcome Trust DBT India Alliance (IA/M/16/1/502606).

## Author contributions

KG, VJ, SC: investigation, original draft preparation; SB, data analysis and original draft preparation; SS and DM: investigation; RP and SSV: conceptualization, supervision, writing-reviewing and editing, funding acquisition.

## Data availability

All original data from the RNA-seq has been deposited in ArrayExpress with accession E-MTAB-9081.

---

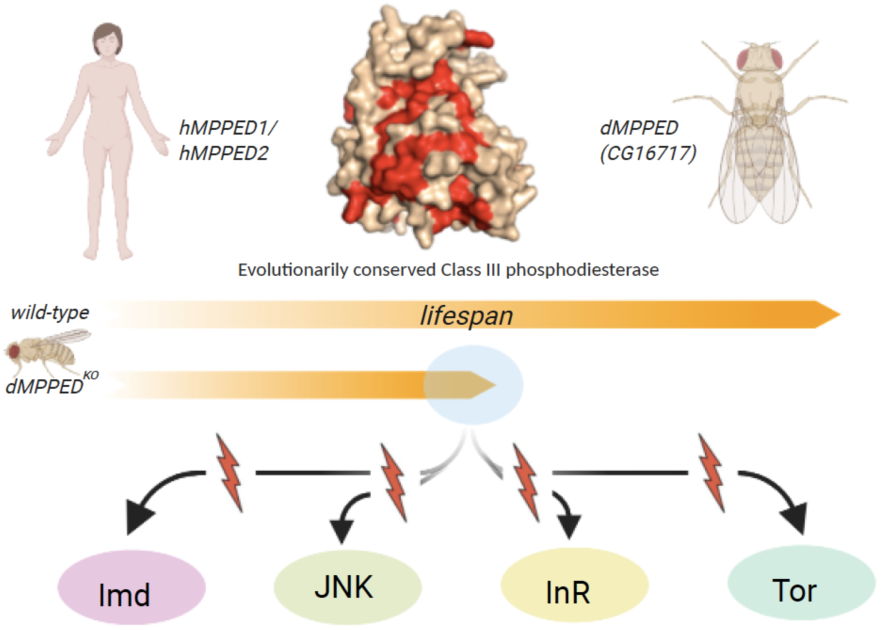
A gene product conserved from the fly to humans is involved in maintaining lifespan in the fly. This activity is not dependent on its metallophosphoesterase activity, and deletion of this gene affects a number of pathways associated with aging in the fly.

