## Supplemental Figure for "An evolutionarily conserved metallophosphodiesterase is a determinant of lifespan in *Drosophila*"

Supplemental Figure 1

**Biochemical characterization of dMPPED**

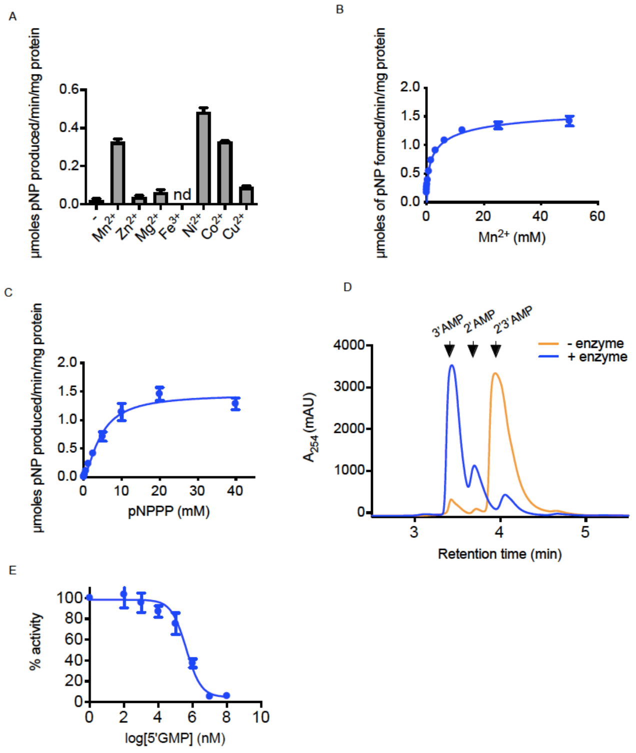


(A**)** Activity of dMPPED (500 ng) was assayed in the presence of 10 mM pNPPP and 5 mM of the indicated metals. Values represent the mean ± S.D. of duplicate determinations carried out using two independent protein preparations. (B) dMPPED (500 ng) was assayed in the presence of 30mM pNPPP and varying concentrations of $Mn^{2+}$. Values represent the mean ± S.D. of duplicate determinations, carried out using two independent protein preparations. (C) dMPPED (500 ng) was assayed in the presence of 5 mM $Mn^{2+}$ and indicated concentrations of pNPPP. Values represent the mean ± S.D. of duplicate determinations using two independent protein preparations. (D) Assays were performed with dMPPED (5 mg) in the presence of 10 mM 2'3'cAMP and 5 mM $Mn^{2+}$. The reaction mixture was analysed using high performance liquid chromatography (HPLC). The HPLC profiles obtained for the enzyme blank (- enzyme) and dMPPED (+ enzyme) have been superimposed. The graph shown is representative of experiments performed twice with two independent protein preparations. (E) dMPPED (500 ng) was assayed in the presence of 30 mM pNPPP, 5 mM $Mn^{2+}$ and indicated amounts of 5'GMP. Inhibition has been plotted as a percentage of the activity shown by dMPPED in the absence of 5'GMP. Values represent the mean ± S.D. of duplicate determinations carried out using two independent protein preparations.

Supplemental Figure 2

| | Adult Male | | Adult Female | | Larval | |
|---|---|---|---|---|---|---|
| **Tissue** | **FPKM** | **Enrichment** | **FPKM** | **Enrichment** | **FPKM** | **Enrichment** |
| Head | 5.3 ± 0.7 | 1.3 | 4.9 ± 0.6 | 0.9 | | |
| Eye | 4.6 ± 0.1 | 1.1 | 4.7 ± 0.2 | 0.9 | | |
| Brain / CNS | 8.5 ± 0.9 | 2.0 | 8.0 ± 0.7 | 1.5 | 5.2 ± 0.8 | 1.2 |
| Thoracicoabdominal ganglion | 6.4 ± 0.8 | 1.5 | 7.7 ± 0.3 | 1.4 | | |
| Crop | 6.9 ± 0.2 | 1.6 | 7.9 ± 0.2 | 1.5 | | |
| Midgut | 4.8 ± 0.5 | 1.1 | 4.9 ± 0.6 | 0.9 | 4.5 ± 0.8 | 1.0 |
| Hindgut | 2.2 ± 0.1 | 0.5 | 2.6 ± 0.3 | 0.5 | 2.7 ± 1.1 | 0.6 |
| Malpighian Tubules | 4.9 ± 1.8 | 1.2 | 5.5 ± 0.1 | 1.0 | 21 ± 6.1 | 4.9 |
| Fat body | 2.7 ± 1.0 | 0.6 | 2.0 ± 1.2 | 0.4 | 5.0 ± 1.1 | 1.2 |
| Salivary gland | 5.1 ± 0.6 | 1.2 | 4.1 ± 0.4 | 0.8 | 4.5 ± 0.3 | 1.0 |
| Heart | pending | — | pending | — | | |
| Trachea | | | | | 6.3 ± 0.4 | 1.4 |
| Ovary | | | 7.4 ± 0.1 | 1.4 | | |
| Virgin Spermatheca | | | 2.1 ± 1.2 | 0.4 | | |
| Mated Spermatheca | | | 2.2 ± 0.6 | 0.4 | | |
| Testis | 15 ± 2.1 | 3.5 | | | | |
| Accessory glands | 3.8 ± 1.3 | 0.9 | | | | |
| Carcass | 4.2 ± 0.2 | 1.0 | 4.2 ± 0.2 | 0.8 | 6.5 ± 0.4 | 1.5 |
| Rectal pad | 5.0 ± 0.3 | 1.2 | 4.6 ± 0.1 | 0.8 | | |
| Whole body | 4.2 ± 0.6 | | 5.4 ± 0.1 | | 4.3 ± 1.0 | |

Table showing expression of *dMPPED* (*CG16717*) provided by FlyBase2 (http://flyatlas.gla.ac.uk/FlyAtlas2/index.html) depicting expression as seen by RNAseq

Supplemental Figure 3

**Generation and Confirmation of dMPPED$^{KO}$ flies**

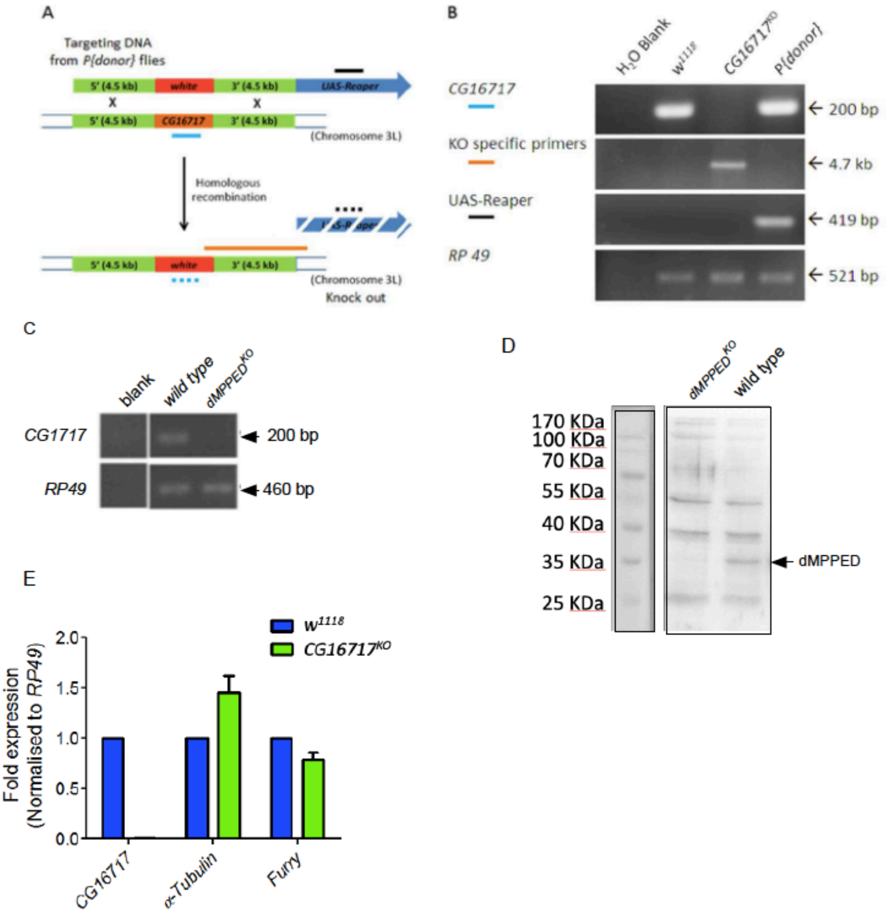


(A) Schematic of homologous recombination at the *dMPPED (CG16717)* locus highlighting the positive selection marker white$^{+}$ and negative selection marker UAS-reaper. Various primers used in subsequent PCRs are depicted as blue, orange or black lines. (B) Genomic PCR of *dMPPED*$^{KO}$ flies using various primers whose amplicons are color coded with the schematic in (A). *Wild type* (*w*$^{1118}$) and P{donor} flies were used as controls for the PCR. (C) RT-PCR performed on *dMPPED*$^{KO}$ flies using *dMPPED* specific primers. *RP49* is used as a loading control. (D) Western blot analysis was performed on protein prepared from brains of flies using dMPPED-specific antibody. The entire protein obtained from 10 brains of indicated genotypes was loaded in each lane. The blot was stripped and re-probed using the anti-tubulin primary antibody as a loading control. (E) Expression levels of *α-Tubulin 67C* and *Furry* in 4 day-old *wild type* and *dMPPED*$^{KO}$ flies. Transcript levels have been normalized to *RP49* and further normalized to the *wild type* flies. Data shown represents mean ± S.D. from 2 independent experiments performed using 10 flies each.
