## Supplemental Table 1 for "An evolutionarily conserved metallophosphodiesterase is a determinant of lifespan in *Drosophila*"

**Summary of all lifespan experiments**

For each lifespan experiment, representative graphs have been presented, and the values in the Table are the average of all experiments. The n values (number of flies assayed), median lifespan, and maximal lifespan (calculated as the median lifespan of the last 10% of the surviving flies) are reported for all the genotypes used in each experiment. The change in median and maximum lifespan is indicated as a percentage compared to the control lines mentioned. The P-value for the log-rank test for these comparisons are indicated. GraphPad Prism 5 was used to plot all the survival curves, calculate the median and maximum lifespan and perform the log-rank test.

| | n | Median lifespan (days) | Maximum lifespan (days) | Change in median lifespan vs control, % | Change in maximum lifespan vs control, % | Log Rank p value | Genotype used to calculate change in lifespan |
|---|---|---|---|---|---|---|---|
| $w^{1118}$ (wild type) | 148 | 66 | 79 | | | | |
| *dMPPED*$^{KO}$ | 132 | 54 | 63 | -18.2 | -20.3 | <0.0001 | $w^{1118}$ |
| *Act-GAL4; dMPPED*$^{KO}$ | 120 | 56 | 59 | | | | |
| *UAS-dMPPED; dMPPED*$^{KO}$ | 127 | 59 | 66 | | | | |
| *Act-GAL4/UAS-dMPPED; dMPPED*$^{KO}$ | 141 | 79 | 87 | 41.1 | 47.5 | <0.0001 | *Act-GAL4; dMPPED*$^{KO}$ |
| $w^{1118}$ (wild type) | 147 | 66 | 79 | | | | |
| *dMPPED*$^{KO}$ | 136 | 54 | 63 | -18.2 | -20.3 | <0.0001 | $w^{1118}$ |
| *elav-GAL4; dMPPED*$^{KO}$ | 149 | 59 | 72 | | | | |
| *UAS-dMPPED; dMPPED*$^{KO}$ | 128 | 59 | 66 | | | | |
| *elav-GAL4;UAS-dMPPED; dMPPED*$^{KO}$ | 140 | 72 | 82 | 22.0 | 13.9 | <0.0001 | *elav-GAL4; dMPPED*$^{KO}$ |
| $w1^{118}$ (wild type) | 208 | 61 | 73.5 | | | | |
| *dMPPED*$^{KO}$ | 275 | 49 | 59 | -19.7 | -19.7 | <0.0001 | $w^{1118}$ |
| *elav-GAL4/+; dMPPED*$^{KO}$ | 230 | 54 | 66 | | | | |
| *UAS-MPPED2*$_{WT}$*/+; dMPPED*$^{KO}$ | 222 | 54 | 75 | | | | |
| *elav-GAL4/UAS-MPPED2*$_{WT}$*; dMPPED*$^{KO}$ | 259 | 66 | 75 | 22.2 | 13.6 | <0.0001 | *elav-GAL4/+; dMPPED*$^{KO}$ |
| $w^{1118}$ (wild type) | 282 | 61 | 66 | | | | |
| *Act-GAL4/+* | 270 | 62 | 76 | | | | |
| *UAS-dMPPED/+* | 286 | 69 | 76 | | | | |
| *Act-GAL4/UAS-dMPPED* | 277 | 76 | 94 | 22.6 | 23.7 | <0.0001 | *Act-GAL4/+* |

| | | | | | | | |
|---|---|---|---|---|---|---|---|
| $w^{1118}$ (wild type) | 142 | 56 | 69 | | | | |
| *elav-GAL4/+* | 134 | 69 | 81 | | | | |
| *UAS-dMPPED/+* | 148 | 69 | 87 | | | | |
| *elav-GAL4/UAS-dMPPED* | 140 | 81 | 93 | 17.4 | 14.8 | <0.0001 | *elav-GAL4/+* |
| $w^{1118}$ (wild type) | 270 | 62 | 78 | | | | |
| *dMPPED*$^{KO}$ | 276 | 52 | 59 | -16.1 | -24.4 | <0.0001 | $w^{1118}$ |
| *elav-GAL4/+; dMPPED*$^{KO}$ | 283 | 62 | 76 | | | | |
| *UAS-dMPPED*$_{D49A}$*/+; dMPPED*$^{KO}$ | 269 | 70 | 89 | | | | |
| *elav-GAL4/UAS-dMPPED*$_{D49A}$*; dMPPED*$^{KO}$ | 265 | 70 | 81 | 12.9 | 6.6 | <0.0001 | *elav-GAL4/+; dMPPED*$^{KO}$ |
| $w^{1118}$ (wild type) | 1118 | 66 | 80 | | | | |
| *elav-GAL4/+* | 148 | 77 | 86 | | | | |
| *UAS-dMPPED*$_{D49A}$*/+* | 148 | 77 | 96 | | | | |
| *elav-GAL4/UAS-dMPPED*$_{D49A}$ | 110 | 85 | 92 | 10.4 | 7.0 | <0.0001 | *elav-GAL4/+* |
| $w^{1118}$ (wild type) | 208 | 61 | 75 | | | | |
| *dMPPED*$^{KO}$ | 275 | 49 | 59 | -19.7 | -21.3 | <0.0001 | $w^{1118}$ |
| *elav-GAL4/+; dMPPED*$^{KO}$ | 230 | 54 | 66 | | | | |
| *UAS-MPPED2*$_{H67R}$*/+; dMPPED*$^{KO}$ | 235 | 54 | 72 | | | | |
| *elav-GAL4/UAS-MPPED2*$_{H67R}$*; dMPPED*$^{KO}$ | 283 | 75 | 81 | 38.9 | 22.7 | <0.0001 | *elav-GAL4/+; dMPPED*$^{KO}$ |
| $w^{1118}$ (wild type) | 1118 | 66 | 80 | | | | |
| *elav-GAL4/+* | 148 | 74 | 86 | | | | |
| *UAS-MPPED2/+* | 84 | 72.5 | 86 | | | | |
| *elav-GAL4/UAS-MPPED2* | 106 | 77 | 86 | 4.1 | 0.0 | 0.2925 (ns) | *elav-GAL4/+* |
