## Supplemental Table 2 for "An evolutionarily conserved metallophosphodiesterase is a determinant of lifespan in *Drosophila*"

Supplemental Table 2 List of primers used in this study

| Name of the primer | Sequence (5’ to 3’) | Annealing Temperature |
|---|---|---|
| dMPPED_D49A_Fwd | GTTTGCATGTCCGCCACGCACTCCCTGAC | |
| RP 49_Fwd | CGGATCGATATGCTAAGCTGT | 60 °C |
| RP 49_Rvs | GCGCTTGTTCGATCCGTA | |
| CG16717_RT_Fwd | ACATGCCGGCGATTTACCAAGTGCG | 60 °C |
| CG16717_RT_Rvs | CGAGGATGGACATGCCGGTGTGTTTG | |
| DILP 2_Fwd | TTTGTCCTTCATCTCGATGGTGGCC | 60 °C |
| DILP 2_Rvs | GCGCTTGTGTGGAATCACGGGATTA | |
| InR_Fwd | AGACGAAATGCCCTGAAAAG | 60 °C |
| InR_Rvs | AAGACACATTTCGACACGAGATG | |
| S6K_Fwd | GACGATGTTGACCTAGAACCGG | 60 °C |
| S6K_Rvs | CACCTTTGCCAAGGACCTTC | |
| 4E-BP_Fwd | ACCACTCCTGGAGGCACCAAACT | 60 °C |
| 4E-BP_Rvs | GGAGTTCCCCTCAGCAAGCAACT | |
| FOXO_Fwd | AGGCTGACCCACACAGATAAC | 60 °C |
| FOXO_Rvs | GGCTCCACAAAGTTTTCGGG | |
| Sirt2_Fwd | ACGCAATTCTATCCGCCAACTAAG | 60 °C |
| Sirt2_Rvs | CGCTTGTTGCTGGTTCTGTGG | |
| Dp110_Fwd | AATCTGCCTGTTGCCCAATG | 60 °C |
| Dp110_Rvs | ATAGCCCAGTGGCATCTGTTT | |
| TotA_Fwd | CTGCTCTTATGTAAGTAGTATCGAAT | 60 °C |
| TotA_Rvs | CAACGATCCTCGCCTTTCGACC | |
| TotM_Fwd | TCGACAGCCTGGTCACTTTC | 60 °C |
| TotM_Rvs | ACCAAGACCACACGAGCATT | |
| Def_Fwd | CGTGGCTATCGCTTTTGCTC | 60 °C |
| Def_Rvs | GAGTAGGTCGCATGTGGCTC | |
| PGRP-SA_Fwd | ACATGCAGGCGTATCATCAGA | 60 °C |
| PGRP-SA_Rvs | CATCCGATGGAAGTTTATCCACA | |
| Dpt_Fwd | ACCGCAGTACCCACTCAATC | 55 °C |
| Dpt_Rvs | CCCAAGTGCTGTCCATATCC | |
| Drs_Fwd | GTACTTGTTCGCCCTCTTCG | 55 °C |
| Drs_Rvs | CTTGCACACACGACGACAG | |
| Dro_Fwd | TTTTCCTGCTGCTTGCTTGC | 55 °C |
| Dro_Rvs | TGATGGCAGCTTGAGTCAGG | |
