## Supplemental Table 3 for "An evolutionarily conserved metallophosphodiesterase is a determinant of lifespan in *Drosophila*"

Supplemental Table 3: List of flies used in this study

| Fly line | Source | Identifier | Microinjection site |
|---|---|---|---|
| *w1118* | BDSC | 3605 | NA |
| *IT-gal41111-G4* | BDSC | 65447 | NA |
| *UAS-mCD8-GFP* | BDSC | 5137 | NA |
| *$foxo^{21}$* | BDSC | 80943 | NA |
| *$foxo^{25}$* | BDSC | 80944 | NA |
| *Act-gal4* | BDSC | 4414 | NA |
| *elav-gal4* | BDSC | 458 | NA |
| P{donor} | This study | This study | Random insertion |
| *dMPPEDKO* | This study | This study | NA |
| *UAS-dMPPED* | This study | This study | *attP40* |
| *UAS-dMPPEDD49A* | This study | This study | *attP40* |
| *UAS-MPPED2* | This study | This study | *attP40* |
| *UAS-MPPED2H67R* | This study | This study | *attP40* |
